# Interleukin-12 elicits a non-canonical response in B16 melanoma cells to enhance survival

**DOI:** 10.1101/608828

**Authors:** C. N. Byrne-Hoffman, W. Deng, O. McGrath, P. Wang, Y. Rojanasakul, D. J. Klinke

**Affiliations:** Department of Pharmaceutical Sciences; West Virginia University, Morgantown, WV 26506; Department of Microbiology, Immunology, and Cell Biology; West Virginia University, Morgantown; WV 26506; Department of Chemical and Biomedical Engineering and WVU Cancer Institute; West Virginia University, Morgantown; WV 26506

**Keywords:** interleukins, intracellular signaling, cytokine, JAK-STAT signaling, PI3K-AKT signaling, non-canonical

## Abstract

Oncogenesis rewires signaling networks to confer a fitness advantage to malignant cells. For instance, the B16F0 melanoma cell model creates a cytokine sink for Interleukin-12 (IL-12) to deprive neighboring cells of this important anti-tumor immune signal. While a cytokine sink provides an indirect fitness advantage, does IL-12 provide an intrinsic advantage to B16F0 cells? Functionally, stimulation with IL-12 enhanced B16F0 cell survival but not normal Melan-A melanocytes that were challenged with a cytotoxic agent. To identify a mechanism, we assayed the phosphorylation of proteins involved in canonical IL-12 signaling, STAT4, and cell survival, Akt. In contrast to T cells that exhibited a canonical response to IL-12 by phosphorylating STAT4, IL-12 stimulation of B16F0 cells predominantly phosphorylated Akt. Mechanistically, the differential response in B16F0 cells is explained by both ligand-dependent and ligand-independent aspects to initiate PI3K-AKT signaling upon IL12RB2 homodimerization. Namely, IL-12 promotes IL12RB2 homodimerization with low affinity and IL12RB2 overexpression promotes homodimerization via molecular crowding on the plasma membrane. Collectively, the data suggest that B16F0 cells shifted the intra-cellular response to IL-12 from engaging immune surveillance to favoring cell survival. Identifying how signaling networks are rewired in model systems of spontaneous origin can inspire therapeutic strategies in humans.

**Statement of Significance:** While Interleukin-12 is a key cytokine that promotes anti-tumor immunity, thinking of cancer as an evolutionary process implies that an immunosuppressive tumor microenvironment could arise during oncogenesis by interfering with endogenous anti-tumor immune signals, like IL-12. Previously, we found that B16F0 cells, a cell line derived from a spontaneous melanoma, interrupts this heterocellular signal by sequestering IL-12, which provides an indirect fitness advantage. Here, we report that B16F0 cells gain an intrinsic advantage by rewiring the canonical response to IL-12 to instead initiate PI3K-AKT signaling, which promotes cell survival. The data suggest a model where overexpressing one component of the IL-12 receptor, IL12RB2, enables melanoma cells to shift the functional response via both IL-12-mediated and molecular crowding-based IL12RB2 homodimerization.

## Introduction

Responses of the different cell types within tissues, such as T cells and Langerhans cells within the epidermal layer of skin, are coordinated by the extracellular release of protein signals. One such extracellular signal is the cytokine Interleukin-12 (IL-12) (1). IL-12 is a heterodimer comprised of p35 and p40 subunits that is produced by antigen presenting cells to boost natural killer (NK) and cytotoxic T lymphocyte (CTL) activity and to polarize T helper cells towards a type 1 phenotype (2). Activated NK, CTLs and type 1 T helper cells play important roles in controlling viral infections and defending against malignancy. Towards this aim, local delivery of IL-12 to the tumor microenvironment has been shown to promote tumor regression in both mouse models and human melanoma (3–8) while knocking out the IL-12 pathway increases malignancy (9).

Inside the cell, these extracellular signals are processed through a series of protein-protein interactions that transmit this information from the cell membrane to the nucleus to orchestrate a cellular response. In the case of IL-12, these extracellular signals are transduced by the canonical Janus-Kinase (JAK)-Signal Transducer and Activator of Transcription (STAT) signaling pathway (10). The IL-12 receptor, composed of a *β*1 (IL12RB1) and *β*2 (IL12RB2) subunit, lacks intrinsic enzymatic activity and forms a complex with two JAKs, JAK2 and TYK2. Autophosphorylation of the receptor complex results in binding and activation of STAT4. Phosphorylated STAT4 dimerizes and then translocates to the nucleus to function as a transcription factor (11). As a transcription factor, STAT4 promotes the expression and release of Interferon-gamma (IFN-*γ*)(12), which upregulates MHC class I antigen presentation (13). An increase in antigen presentation enables CTLs to target and kill virally infected and malignant cells present within the tissue (14).

While most of our understanding of how information is transmitted between cells is based on normal biology, cancer cells arise through a process of mutation and selection (15, 16). This implies that cancer cells develop ways during oncogenesis to block these extracellular signals or rewire intracellular signaling pathways to gain a fitness advantage (17, 18). Of note, B16F0 cells correspond to the parental cell line isolated from a spontaneous tumor that developed in C57BL/6J mice and are used as a syngeneic transplantable mouse model for human melanoma (19, 20). In contrast to genetically engineered mouse models where genetic alterations that promote oncogenic transformation are invoked a priori (21), tumor cell lines derived from spontaneous tumors are likely to have a broader collection of mechanisms that promote oncogenic transformation. Following from this context, the mouse melanoma cell line B16F0 overexpresses one component of the IL-12 receptor, IL12RB2, to form a cytokine sink for IL-12, which can deprive neighboring immune cells of this important anti-tumor cytokine (22). As rewiring of intracellular cell signaling networks is another way that tumor cells can gain advantage, we asked here whether IL-12 provides an intrinsic advantage to B16F0 tumor cells.

## Materials and Methods

### Cell Culture

B16F0 mouse melanoma (ATCC, Manassas, VA), Melan A mouse melanocytes (V. Hearing, National Cancer Institute, Bethesda, MD) and 2D6 cells (M. Grusby, Harvard University, Cambridge, MA) were cultured as described previously (23). All cell lines were revived from frozen stock and used within 10-15 passages that did not exceed a period of 6 months.

### Apoptosis and Necrosis Assay

Cells were plated in a 12-well tissue culture plate (BD Falcon, BD Biosciences, San Jose, CA) for 12 hours, then pre-treated with complete culture media with a final concentration of 40, 100, or 200 ng/ml IL-12. After 18 hours of pretreatment, cells were treated with 0, 5, 10, or 20 *µ*M Imatinib (Cayman Chemical, Ann Arbor Michigan). A 0.1% DMSO vehicle control in complete media was used as a negative control, while cells heated to 50*°*C for 5-10 minutes or exposed to 10% DMSO served as positive controls. After 24 hours of incubation with imatinib or other treatment, cell viability was assessed using either a flow cytometry-based assay or an ATPlite assay (PerkinElmer, Shelton, CT), which was performed according to the manufacturer’s instructions. For the flow cytometry-based assay, cell media and floating cells were pooled with cells detached with DPBS without magnesium or calcium, centrifuged in 15 mL conical tubes at 250xg, and cells transferred to a 96 well round bottom plate (BD Falcon) for staining. Cells were then washed with DPBS with magnesium and calcium. After washing, cells were stained with an apoptosis and necrosis staining kit (ABD Serotech, Bio-Rad, Raleigh, NC) according to manufacturer’s instructions and analyzed by flow cytometry on a BD LSRFortessa. Data was analyzed in the BD FACSDiva 3.0 software and in Microsoft Excel. Statistical significance in the difference in viability upon IL-12 treatment for a given imatinib concentration was assessed using a two-tailed homoscedastic t-Test, where a p-value less than 0.05 was considered significant.

### Flow Cytometry

Cells were grown in a 12-well tissue culture plate if adherent (B16F0 and Melan-A) or in a 96-well round bottom plate if non-adherent (2D6) (BD Falcon). At each time point, adherent cells were detached from the plate with Trypsin-EDTA or DPBS without calcium/magnesium. For IL12RB1 and IL12RB2 copy number, cells were stained for 30 minutes with Yellow or Violet Live/Dead Fixable Stain (BD Biosciences), fixed with Lyse/Fix Buffer (BD Biosciences), then stained for PE-IL12RB1 (BD Biosciences) and AlexaFluor 488-IL12RB2 (R&D Systems, Minneapolis, MN) antibodies. Quantum Simply Cellular microspheres (Bangs Laboratories, Fishers, IN) were stained with IL12RB1 or IL12RB2 antibody within the same experiment to assess receptor copy number. Single stained controls were used to assess fluorescent spillover. To assay STAT4 and Akt activity, B16F0 and 2D6 cells were starved of FBS and IL-12p70 for 12-24 hours and then treated for 1hr or 3hr with the indicated concentration of IL-12p70 or vehicle. Positive control cells were treated with 10% FBS and 40 ng/mL IL-12p70. Cells were stained using Violet Fixable Live/Dead (Invitrogen) to assess cell viability, fixed (BD Biosciences Lyse/Fix), permeabilized with Perm Buffer III (BD Biosciences), blocked with Mouse IgG (1:100, concentration, Jackson Labs, West Grove, PA), stained with FITC-IL12RB2 (R&D Systems), PE-pAKT and AF647-pSTAT4 (BD Biosciences) and stored at 4*°*C in PBSaz before analysis. Cells were analyzed using the BD LSRFortessa flow cytometry analyzer, FCSExpress, and BD FACSDiva 3.0 software. Following acquiring 20,000 events on average, forward and side scatter were used to gate cell populations, where live cells were identified using Live-Dead staining and used for subsequent analysis.

### CRISPR/Cas9 Constructs, Transfection and Cloning Validation

Two vectors were prepared by GenScript (Piscataway, NJ) with puromycin resistance genes and the flanking guide RNA (gRNA) sequences TCCGCATACGTTCACGTTCT and AGAA-TTTCCAAGATCGTTGA and transfected into wild-type B16F0 cells using Lipofectamine 2000 (Thermo Fisher Scientific, Waltham, MA). Transfection was confirmed via microscopy visually using a green fluorescent protein control. Derivatives of the B16F0 cell line were obtained from single clones isolated using puromycin selection and single cell plating. About 25% of clones exhibited some level of IL12RB2 knockdown, which was confirmed by flow cytometry.

### Western Blot

B16F0 and 2D6 cells were grown in complete media in 6-well plates 24 hours before the beginning of the experiment. At 12 hours prior, cells were starved of FBS and IL-12. One hour before start of experiment, cells were treated with control media, PI3K inhibitor LY294002 (50 *µ*M, Cell Signaling Technology, Danvers, MA), or with Akt inhibitor MK-2206 (2.4 ug/mL or 5 *µ*Mol/L, Cayman Chemical, Ann Arbor, MI). Cells were then treated with 100 ng/mL IL-12 at t=0 or stimulated with FBS as a positive control. After 30 minutes or 3 hours, cells were collected for cell lysates for western blot analysis for total Akt (1:1000, Cell Signaling Technology), pAkt (1:1000, Cell Signaling Technology), and *β*-actin (1:1000, Sigma Aldrich, St. Louis, MO). The cells were lysed in radioimmunoprecipitation assay (RIPA) buffer (50 mM Tris-HCl, pH 8.0, 150 mM NaCl, 1% Nonidet P-40, 0.5% sodium deoxycholate, 0.1% SDS, 2 mM EDTA, 1 mM NaVO4, 10 mM NaF, and protease inhibitors), and the protein concentration in the lysates was determined by a spectrophotometer. Equal amounts of the lysates were subjected to SDS-polyacrylamide gel electrophoresis (PAGE) and the immunoblotting was performed with use of the primary antibodies listed above and secondary antibodies conjugated to horseradish peroxidase (Jackson ImmunoResearch, West Grove, PA). Proteins were visualized by chemiluminescence.

### Data Analysis and Statistics

Single-cell gene expression (scRNA-seq) analysis of 31 human melanoma tumors were downloaded from the Gene Expression Omnibus (GEO) entry GSE115978 (24). All data was analyzed in R (V3.5.1) using the ‘stats’ package (V3.5.1). A Pearson’s Chi-squared test was used to assess statistical differences between distributions.

The impact of IL-12 on decreasing the cytotoxic efficacy of imatinib was inferred from the data using a mathematical model:

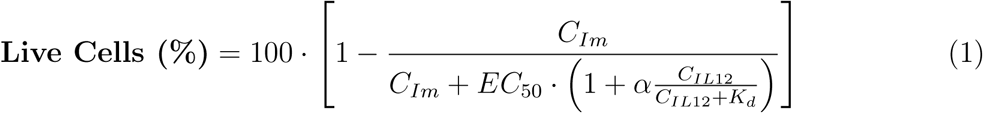

where *C_Im_* is the concentration of imatinib, *C_IL_*_12_ is the concentration of IL-12, *EC*_50_ is the effective concentration required to elicit a 50% drop in viability, *K_d_* is the equilibrium binding constant of IL-12 and set equal to 40 ng/mL, and *α* is a parameter that quantifies the impact of IL-12 in shifting the *EC*_50_ of imatinib. As described previously (25), a Markov Chain Monte Carlo approach based on a Metropolis-Hasting algorithm was used to obtain converged Markov Chains that represent samples from the distributions in the model parameters that were consistent with the data, that is posterior distributions in the model parameters. In brief, three Markov Chains were generated with over-disperse starting points. New steps in the chains were proposed using a normally distributed random number generator with a mean of zero and adjusted standard deviation such that the acceptance fraction was 0.2. The Gelman-Rubin potential scale reduction factor applied to the parameters was used to assess convergence of the chains to sampling of the posterior distribution. Performance and diagnostic metrics of the chains are shown as supplementary Figures S2 and S4. The posterior distribution in parameter *α* was used to infer whether the effect was greater than zero and significant. A p-value (*P* (*α <* 0*|Model, data*)) of less 0.05 was considered as significant, that is the data support that IL-12 provides a rescue effect to imatinib. The significance associated with differences in STAT4 and Akt phosphorylation was assessed using a two-tailed homoscedastic t-Test, where a difference with a p-value less than 0.05 was considered significant.

### Mathematical Modeling and Model Selection

Generally, the IL-12 signaling pathway transduces signals by activating Akt and STAT4 in three steps. Here, we postulate that the first step is one of three competing mechanisms that create an activated receptor complex: a canonical ligand-dependent mechanism, a non-canonical receptor-dependent mechanism, and a hybrid model comprised of both canonical and non-canonical mechanisms. The canonical ligand-dependent mechanism corresponds to the reversible binding of a ligand (*IL*12) to a receptor multi-protein complex (*RC*) resulting in a conformation change that increases kinase activity of the complex (*ARC*):

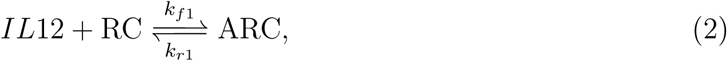

with the forward and reverse rate constants *k_f_* _1_ and *k_r_*_1_, respectively. Assuming that the total number of receptor complexes (*T OT RC*) are conserved, the amount of activated receptor in equilibrium with the ligand concentration is given by:

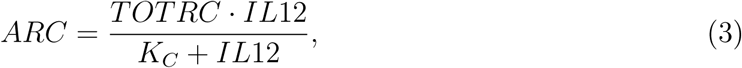

where *K_C_* is equal to *k_r_*_1_*/k_f_* _1_. Similarly, a non-canonical receptor-dependent mechanism represents the spontaneous change in activity of a receptor complex:

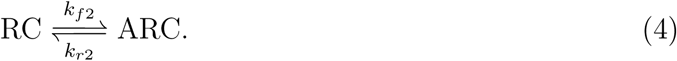

At equilibrium, the amount of activated receptor is given by:

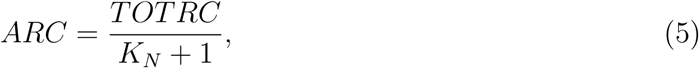

where *K_N_* is equal to *k_r_*_2_*/k_f_* _2_. The hybrid model combines both mechanisms:

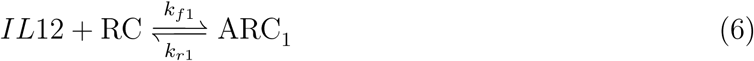

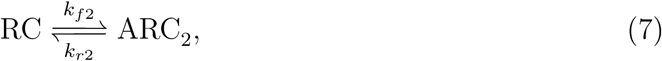

with the equilibrium relationships:

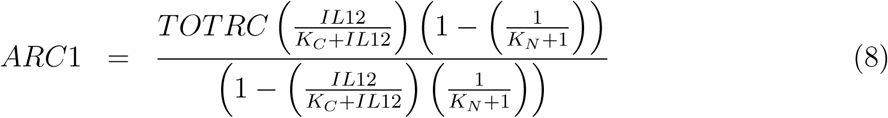

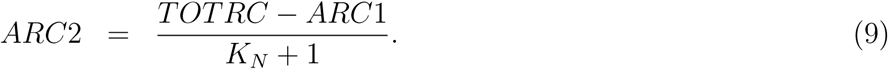

Based on the relative abundance of IL-12 receptor proteins, we assumed that the total receptor complex (*T OT RC*) is comprised primarily of IL12RB1 and IL12RB2 heterodimers in 2D6 cells while IL12RB2 homodimers are primarily present in B16F0 cells and correspond to the abundances measured by flow cytometry. The concentration of IL12 is an experimental variable and the parameters *K_C_* and *K_N_* were fit to the experimental data.

An activated receptor complex catalyzes the second step by phosphorylating (*p−*) the signaling intermediate (*S*), such as Akt or STAT4:

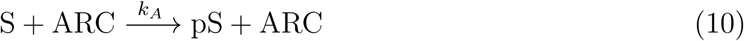

with the rate constant for phosphorylation, *k_A_*. While two different mechanisms activate the receptor complex (e.g., ARC1 versus ARC2), we assumed that the phosphorylation kinetics of the signaling intermediate are the same. Activation is coupled with a deactivation step:

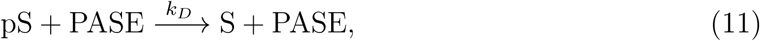

which is catalyzed by a generic phosphatase (*PASE*) with a rate constant of *k_D_* and captures the decline in phosphorylation upon removal of the ligand. Assuming the total signaling intermediate is conserved (*T OT S*), the amount of phosphorylated signaling intermediate for a given abundance of activated receptor complex (*ARC*) at steady state is:

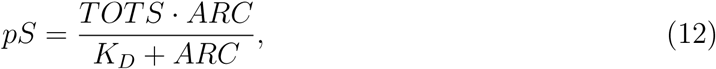

where *K_D_* is equal to *k_D_ ·PASE/k_A_*. *T OT S* and *K_D_* were free parameters used in calibrating the mathematical models to the data. The three competing models are expressed as:

**Canonical:**

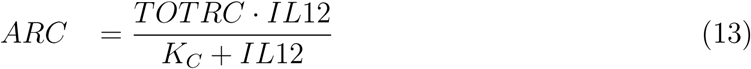

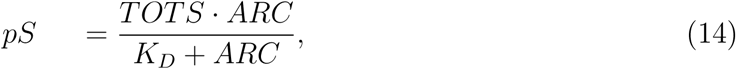

**Non-canonical:**

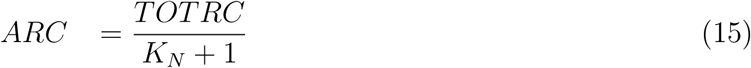

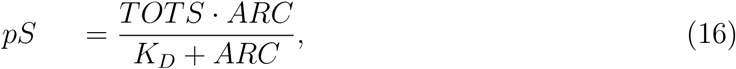

**Hybrid:**

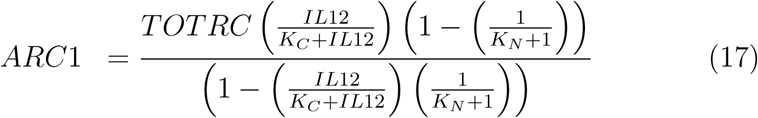

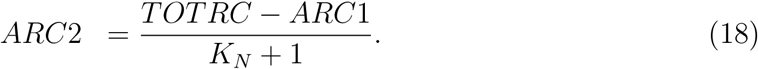

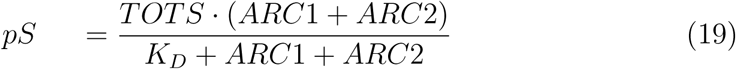

Except for *K_C_*, the three mathematical models were fit separately to Akt and STAT4 phosphorylation (expressed in MFI) versus IL12RB2 data (expressed in copies) for each cell line stratified by IL12RB2 expression and observed under unstimulated and IL-12 stimulated conditions. The ligand binding constant, *K_C_*, was determined for each cell line, as extracellular binding affinity should not influence the specificity of intracellular signal transduction. As the number of experimental observations (*N_obs_ ≥* 69) exceeded the number of free parameters (*N_parameters_ ≤* 4), the model parameters were theoretically identifiable from the data. A summed squares of the error (SSE) metric was used as a goodness-of-fit for model *i*:

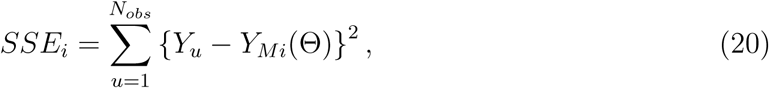

which quantifies the deviation between specific observations, *Y_u_*, and their respective prediction by model *i*: *Y_Mi_*. The maximum likelihood estimate for the parameters (Θ*_ML_*) of the corresponding mathematical models were obtained at the minimum SSE. To quantify the strength of evidence favoring model *i* over model *j*, we used the Bayes Factor (25):

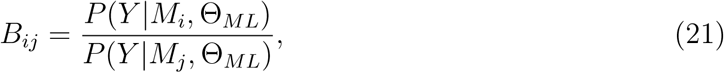

where *P* (*Y |M_i_,* Θ*_ML_*) is the likelihood for observing data (*Y*), given the mathematical model *i* and parameter values Θ*_ML_*, and can be estimated by:

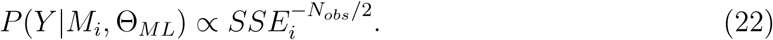

where *N_obs_* is the number of observations of the experimental variable. To interpret the Bayes Factor, values for the *B_ij_* between 1 and 3 are considered weak, between 3 and 20 are positive, between 20 and 150 are strong, and *>*150 are very strong evidence favoring model *i* over model *j* (26).

## Results

To investigate whether IL12RB2 provides an intrinsic advantage to the B16F0 cell line, we first asked whether the subunits of the IL-12 receptor are similarly upregulated in Melan-A cells, a model of ‘normal’ melanocytes. The 2D6 T cells, a mouse type 1 T helper cell line, were used as a positive control for IL-12 receptor expression (22). IL12RB1 and IL12RB2 abundance was assayed by flow cytometry and calibrated to copy number using antibody calibration beads (Figure 1).

**Figure 1:**
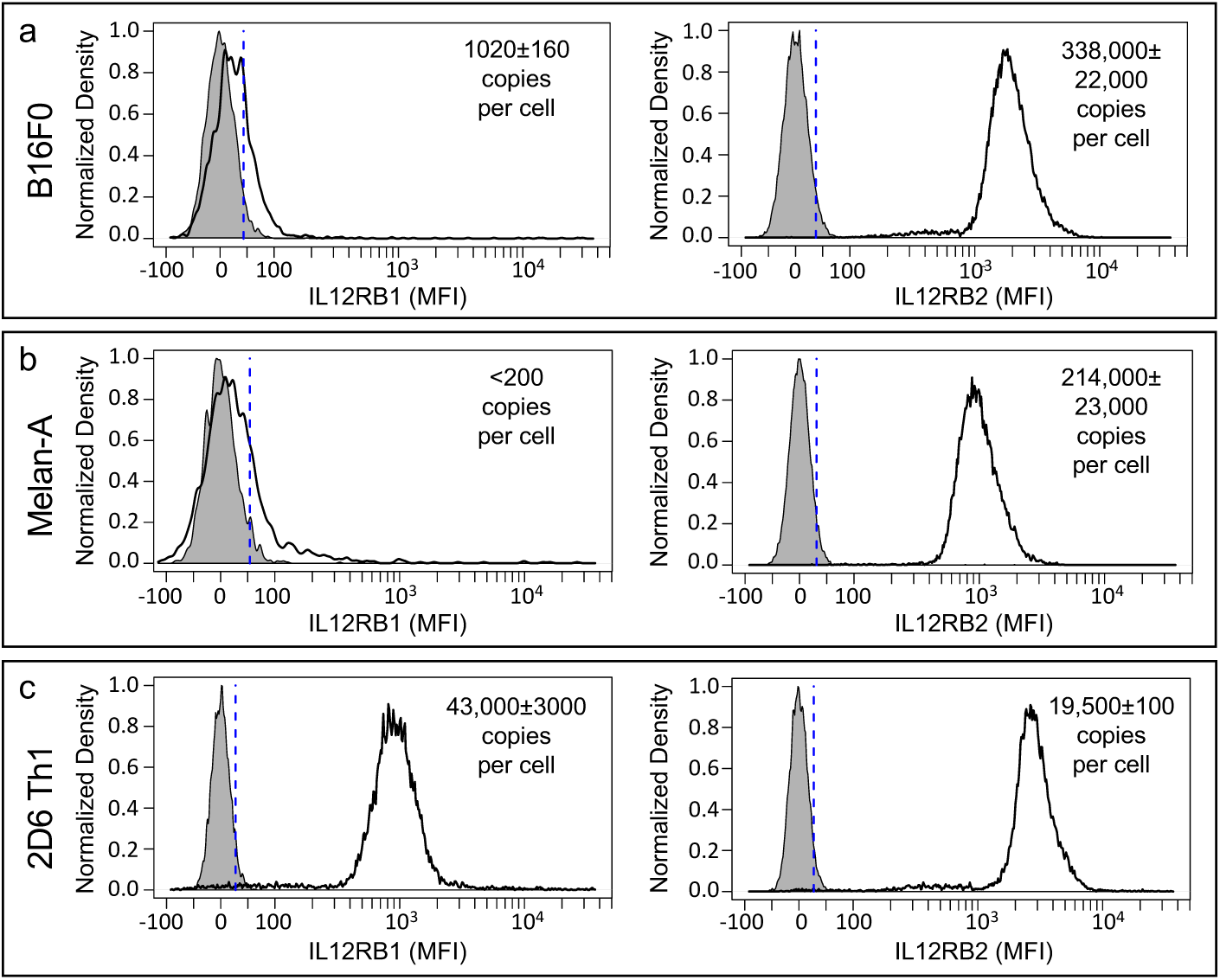
Components of the IL-12 receptor are differentially expressed in mouse melanoma cell lines compared to T cells. IL12RB1 (left panels) and IL12RB2 (right panels) expression in B16F0 melanoma (a), Melan-A ‘normal’ melanocytes (b), and 2D6 Th1 T helper cells (c), were assayed by flow cytometry. Antibody-binding calibration beads were used to determine copy numbers. No stain samples were used as negative controls (gray shaded curves). Results representative of at least three biological replicates.

In 2D6 cells, IL12RB1 and IL12RB2 were present in almost a 2:1 copy ratio, with 43,000 copies of IL12RB1 and 19,500 copies of IL12RB2 and consistent with previous findings (22). Interestingly, the melanoma-related cell lines stained brightly for IL12RB2. The copy number of IL12RB2 was 1.5 times higher in the melanoma cell line versus the ‘normal’ melanocyte line, where the B16F0 cell line expressed over 330,000 copies of IL12RB2 and Melan-A cells had 214,000 copies. In contrast, IL12RB1 abundance was barely detectable in both cell lines, as staining was similar to the unstained controls. Only the B16F0 cell line was quantified at 1000 copies, while the Melan-A cell line was below the detection level of 200 copies per cell. These data suggest that expression of components of the IL-12 receptor, IL12RB1 and IL12RB2, are differentially regulated in mouse cell models of melanoma compared with T cells. This differential regulation is manifested as a high copy number of IL12RB2 and an absence of IL12RB1, such that copies of the heterodimeric IL-12 receptor are limited. Of note, we previously reported that the apparent molecular weight of IL12RB2 is similar between B16F0 and 2D6 cells when assayed by Western blot using an antibody recognizing the C-terminus (see Figure 1F in (23)). As the antibody used for flow cytometry recognizes the extracellular domain of IL12RB2 (N-terminus), the Western blot and flow cytometry results suggest that a similar isoform of IL12RB2 is expressed in B16F0 and 2D6 cells.

As B16F0 cells do not make IL-12 (see Figures 2D and S2A in (22)), creating a cytokine sink by overexpressing IL12RB2 provides an indirect survival advantage to malignant melanocytes. We also wondered whether overexpression of IL12RB2 provides a direct survival advantage to malignant melanocytes. To test for an intrinsic survival advantage, we used an apoptosis and necrosis assay to assess cell viability during a chemotherapy challenge with imatinib in the B16F0 melanoma (Figure 2) in combination with an IL-12 rescue. We used imatinib, as it is being investigated as a possible chemotherapy for metastatic melanoma (27, 28). To determine whether IL-12 could rescue the melanoma cells from the cytotoxic effects of chemotherapy, we incubated the cells with three different concentration of IL-12 (40, 100, or 200 ng/mL) for 18 hours, which correspond to concentrations of IL-12 in the serum of various mouse models at approximately 3 – 134 ng/mL (29). We then treated with three different concentrations of imatinib (5, 10, or 20 *µ*M) for 24 hours, with 0.1% DMSO in DPBS serving as a vehicle control. Cell viability was assayed by flow cytometry, whereby cells were identified based on forward and side scatter characteristics. Gating on “cells”, the presence of Annexin-V staining on the cell surface and no PI staining indicates a cell undergoing apoptosis, while cells that are positive for PI and negative for Annexin-V indicates a cell that underwent necrotic cell death.

**Figure 2:**
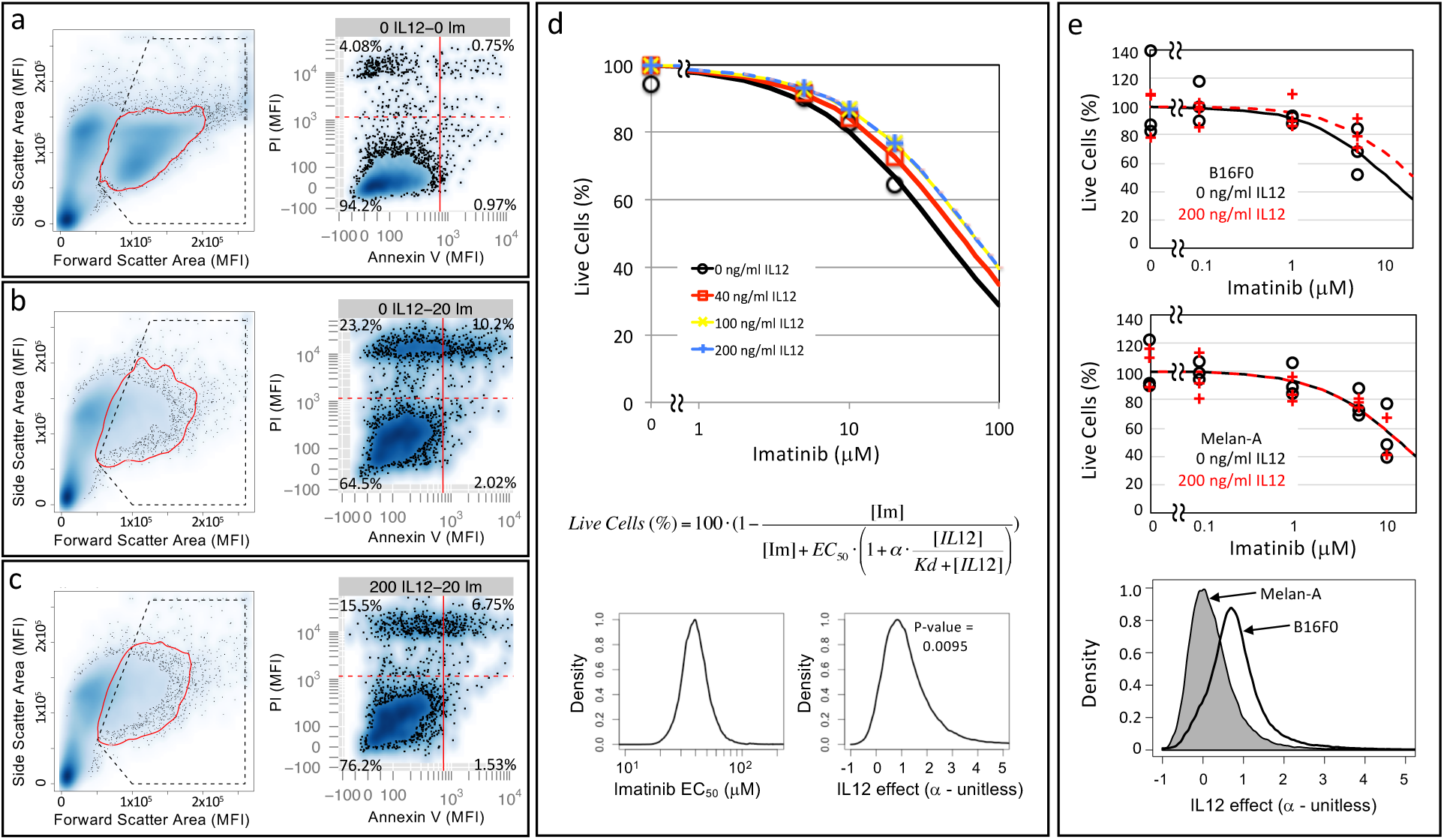
B16F0 melanoma cells but not Melan-A melanocytes can be rescued from the cytotoxic effect of imatinib by IL-12. Cell viability in response to different concentrations of IL-12 and imatinib was assayed by flow cytometry, whereby representative results for specific concentrations (a: No IL-12, No imatinib; b: 20 *µ*M imatinib; c: 200 ng/mL IL-12 and 20 *µ*M imatinib) are shown. Cell events were identified based on forward and side scatter properties (enclosed in red curve in left panel). Annexin V and PI staining were used to identify viable cells (PI negative, Annexin V negative), cells undergoing apoptotic (Annexin V positive, PI negative) and necrotic (Annexin V negative, PI positive) cell death, and cells in late stages of apoptosis (Annexin V positive, PI positive). (d) Using the collective observed data (symbols), a mathematical model was used to estimate the increase in *EC*_50_ of imatinib in the presence of IL-12. The curves trace the corresponding maximum likelihood predictions for each experimental condition. A Markov Chain Monte Carlo approach was used to estimate the model parameters. The posterior distributions in the *EC*_50_ of imatinib and the rescue effect of IL-12 (*α*) are shown in the bottom panels. (e) Viability of B16F0 cells (top panel) and Melan-A cells (middle panel) were assessed using an ATPlite assay following exposure to the indicated concentrations of imatinib in the presence (red curves) or absence (black curves) of 200 ng/mL IL-12. Results for three biological replicates at each condition are shown. The resulting data (symbols) were analyzed using a mathematical model, where the maximum likelihood predictions for each experimental condition are indicated by the curves. Using a Markov Chain Monte Carlo approach, the posterior distributions in the rescue effect of IL-12 (*α*) are shown in the bottom panels (Melan-A: grey shaded, B16F0: black curve).

As summarized in Figure 2, the viability of B16F0 cells was greater than 90% at baseline. Upon treatment with imatinib, the viability was decreased in a dose-dependent manner, as expected. At 24 hours, very few of the cells were observed during the early stages of apoptosis (Annexin-V+, PI-) and the majority of non-viable cells either underwent necrotic cell death (Annexin-V-, PI+) or were in the late stages of apoptotic cell death (Annexin-V+, PI+). The addition of IL-12 reduced the cytotoxic efficacy of imatinib such that the maximum shift in viability of cells pre-treated with IL-12 before imatinib challenge was about 15%. To quantify the effects of imatinib and IL-12 on the viability of the B16F0 cells, we analyzed the data using a mathematical model and inferred the model parameters using a Markov Chain Monte Carlo (MCMC) approach. In theory, the model is identifiable as there are more data points (n=16) than model parameters: *EC*_50_ and *α*. However given the non-linear nature of the model, the MCMC approach was used to assess whether the model parameters could be practically identified from the available data in addition to estimating parameter values. Traces from three independent chains initiated with different starting values and the corresponding Gelman-Rubin convergence diagnostics are shown in Figure S2. Using the converged segments of the chains, the MCMC results suggested that the effective concentration of imatinib (*EC*_50_) and the rescue effect of IL-12 could be practically identified from the data as they exhibited symmetric bounded distributions. The posterior distributions provided a median value of the *EC*_50_ of imatinib equal to 40 nM for imatinib and of *α* equal to 1 (Figure 2d). Moreover, the posterior distributions indicated that the rescue effect of IL-12 from the cytotoxic actions of imatinib was significant (p = 0.0095 that *α <* 0). Quantitatively, a saturating concentration of IL-12 would increase the *EC*_50_ for imatinib by 100% (maximum likelihood value of *α*=1), thereby shifting the imatinib dose-response curve to the right. We repeated this experiment using a plate-based viability assay (ATPlite) using both B16F0 and Melan-A cells (Figure 2e). We used a calibration curve to confirm that these results were within the dynamic range of the assay (see Figure S3a). Of note, higher concentrations of imatinib resulted in near complete cell death, although the results were outside of the dynamic range and thus not quantitative (see Figure S3b). While the variability of the ATPlite assay was higher than the flow cytometry-based assay, IL-12 did not provide a rescue effect when Melan-A cells, an immortalized normal melanocyte cell line, were exposed to imatinib. Specifically, MCMC-based analysis of the ATPlite-generated data with Melan-A cells provided a posterior distribution in *α* with a maximum likelihood value of 0 (see bottom panel of Figure 2e and Figure S4 for MCMC diagnostics). While cell viability was assayed after only 24 hours, the observed difference observed with B16F0 cells implies that, over a 1 week period, tumor load would increase by about 10-fold in the IL-12 pre-treated cells versus chemotherapy alone.

To confirm the role of IL12RB2 in the rescue effect of IL-12 to cytotoxic stress with imatinib, we knocked down IL12RB2 in the B16F0 cell line using a CRISPR-Cas9 construct (Figure 3a). Generally, knockdown clones grew at similar rates, showed minor differences in color, and displayed similar morphology to the wild-type cell line. Knockdown of IL12RB2 was confirmed by flow cytometry. To compare the functional response, wild type (wt) and IL12RB2 knockdown (KD) B16F0 cells were treated with increasing concentrations of imatinib for 12 hours and pre-stimulation with 100 ng/mL IL-12 overnight was included as a second independent variable. Cell viability was assessed using an ATPlite assay. Consistent with data shown in Figure 2, imatinib dose-dependently reduced viability in the wild-type B16F0 cell line that was prevented with pre-treatment with IL-12 (Figure 3b). The rescue effect of IL-12 was most pronounced at the higher concentrations of imatinib. In comparison, imatinib also dose-dependently reduced viability of the IL12RB2 knockdown cells but IL-12 provided no benefit (Figure 3c). Visually, we also noticed that IL-12 had no effect on preventing cell detachment or inhibiting growth in IL12RB2 KD cells. Collectively, the results suggested that IL-12 enhances the survival of B16F0 cells in the presence of cytotoxic stressors, like imatinib, and its effect is dependent on IL12RB2 expression.

**Figure 3:**
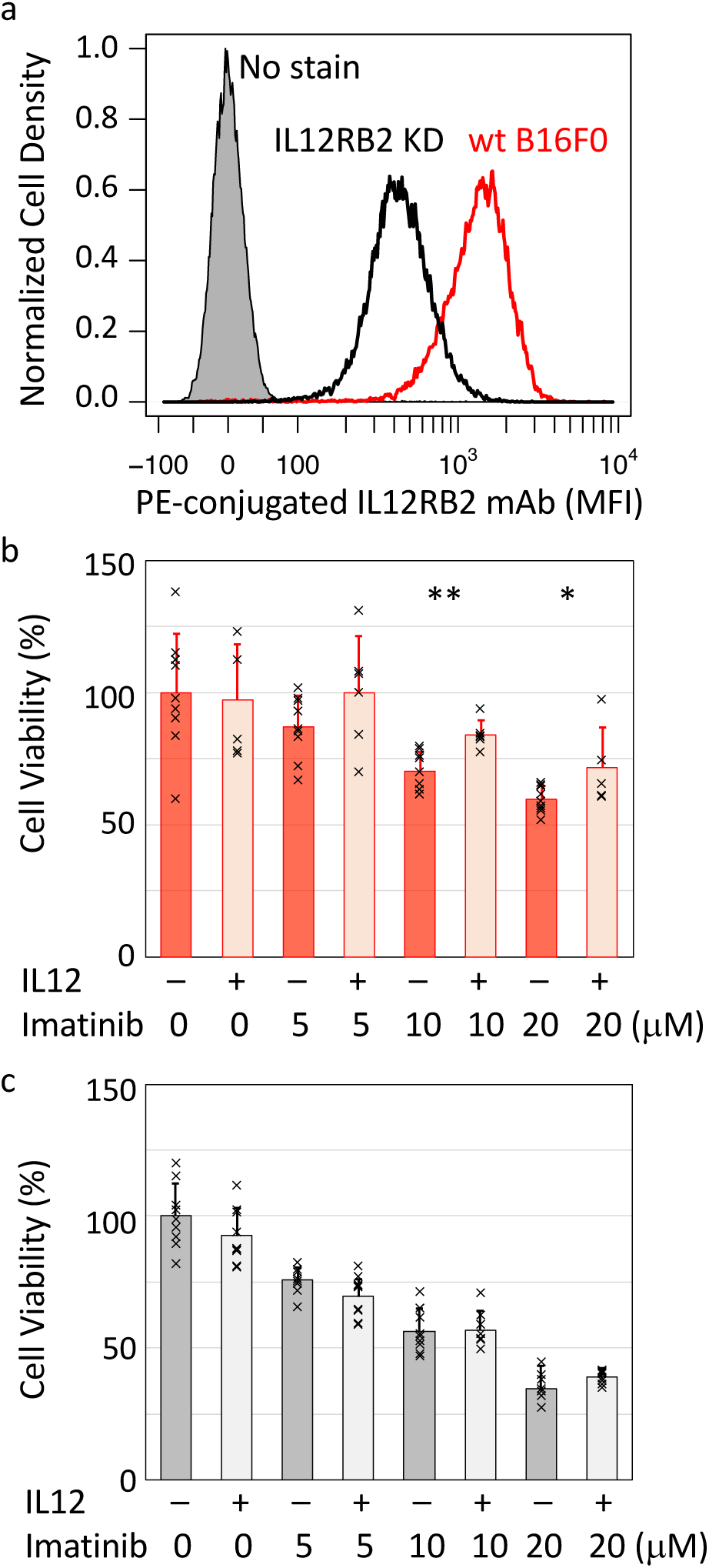
IL12RB2 knockdown in B16F0 decreases the survival benefit of IL-12. (a) Flow cytometry was used to compare IL12RB2 expression in B16F0 cells edited using a CRISPR/Cas9 construct to knockdown IL12RB2 (IL12RB2 KD – black histogram) with wild-type (wt – red histogram) B16F0 cells. Unstained B16F0 cells were used as a negative control (grey shaded histogram). Cell viability was assessed using an ATPlite assay in wild-type (b) and IL12RB2 KD (c) B16F0 cells following treatment with the indicated concentration of imatinib with or without pre-treatment with 100 ng/ml of IL-12. Results are summarized by a bar graph (ave +/- stdev) overlaid with individual results (x’s) that represent at least 5 independent biological replicates for each condition. Statistical significance in the difference in viability upon IL-12 treatment for a given imatinib concentration was assessed using a two-tailed homoscedastic t-Test, where ** indicates a p-value *<* 0.005 and * indicates a p-value *<* 0.05.

Given the imbalance of IL-12 receptor expression in B16F0 cells, it was unclear what signals were transduced in B16F0 cells in response to IL-12 stimulation. Moreover, activating STAT4, which plays a role in Interferon-*γ* gene expression and not survival, is thought to be the canonical signaling response to IL-12. To investigate, we assayed the phosphorylation of STAT4, a canonical signal transducer, and Akt, a non-canonical signal transducer involved in promoting cell survival and therapeutic resistance (30), in response to IL-12 stimulation in both live B16F0 and 2D6 cells by flow cytometry (Figure 4). The 2D6 T helper cell line was used as a model for canonical IL-12 signal transduction. Following 12-24 hours of preconditioning in media free of IL-12 (2D6 only) and serum (2D6 and B16), we stimulated B16F0 and 2D6 cells with increasing concentrations of IL-12 (0, 7, 100, and 300 ng/ml). Pre-conditioning ensured that activation of the IL-12 signaling pathway was due to IL-12, and not due to factors in the serum or basal IL-12. PBS was used as a negative vehicle control for both cell lines.

**Figure 4:**
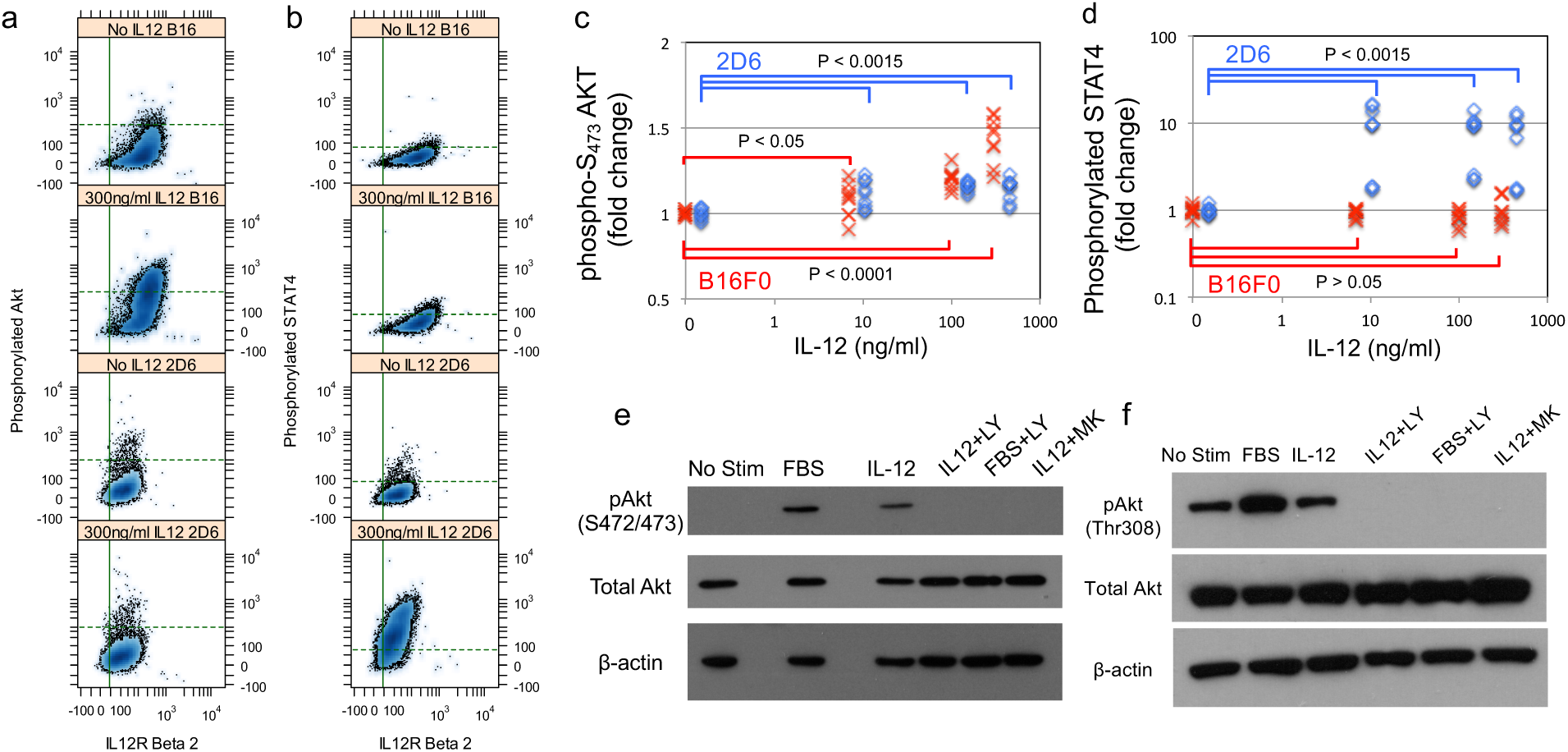
B16F0 melanoma cells transduce non-canonical signals in response to IL-12 stimulation by phosphorylating Akt on serine 472/473. Following a 12-hour preconditioning in the absence of FBS and IL12, B16F0 (top panels) and 2D6 (bottom panels) cells were stimulated with 7, 100, and 300 ng/mL IL-12 for 3 hours and assayed for phosphorylation of Akt (a) and STAT4 (b) by flow cytometry. Background levels of fluorescence associated with phosphorylated Akt and STAT4 in unstimulated cells are represented by the horizontal dotted lines (95% of population below), while the vertical lines indicate background fluorescence associated with IL12RB2 expression in no stain controls (95% of population with lower values). Results for three independent experiments with 3 biological replicates in each are summarized in panels (c) and (d), with p-values for pair-wise comparisons indicated. Activation of Akt by IL-12 (100 ng/mL) in B16F0 was probed using antibodies that recognized phosphorylation at serine 472/273 (e) and threonine 308 (f). IL-12-stimulated cells were compared against unstimulated (negative control), 10% FBS stimulated (positive control), and cells stimulated with IL-12 plus PI3K inhibitor LY294002 (IL12+LY), FBS plus PI3K inhibitor (LY294002), or IL-12 plus pAkt selective inhibitor (IL12+MK). Beta-actin and total Akt serve as loading controls. Results are representative of at least 3 biological replicates.

Only 2D6 cells phosphorylated STAT4 in response to IL-12 stimulation (p*<*0.0015 for 2D6 and p*>*0.05 for the B16), as expected. Basal STAT4 phosphorylation was minimal in both cell lines for the vehicle only-treated controls. In terms of Akt phosphorylation, the B16 melanoma showed a marked activation of Akt within 3 hours at all concentrations of IL-12 tested, with a progressive increase in pAkt with increasing concentrations of IL-12 treatment (from p *<* 0.05 to p *<* 0.0001). In contrast, 2D6 cells exhibited a less pronounced shift in phosphorylation of Akt above baseline in response to IL-12 treatment (p*<*0.0015). Pre-conditioning cells in the absence of serum was critical to observe differences in Akt phosphorylation.

To determine whether Akt activation in the B16 melanoma line was specific, we repeated the experiment and used a Western Blot assay to determine basal levels of Akt and test several inhibitors against the IL-12-induced effect on Akt phosphorylation (Figure 4 panels e and f). To serve as a negative control and to determine whether the PI3K pathway may be cross-activated in response to IL-12 stimulation, we used a PI3K inhibitor LY294002. In addition, we used a selective pAkt inhibitor, MK-2206, to investigate whether the observed effect was solely through possible direct activation of Akt by the IL-12 receptor. Two different phosphorylation sites on Akt were assessed, threonine 308 (Thr308) and serine 472/473 (Ser472/473). We also measured total Akt, and *β*-actin was used as a loading control. We found no change in phosphorylation of Thr308 in response to IL-12 stimulation, whereas baseline phosphorylation was turned off by both LY294002 and MK-2206. However, IL-12 stimulation produced a marked increase in phosphorylation of the Ser472/473 site, and again, phosphorylation was blocked by both the PI3K and Akt-selective inhibitors.

Conventionally, signal transduction within the IL-12 pathway is comprised of three steps. The first step involves a change in conformation of the multi-protein receptor complex upon ligand binding that promotes receptor kinase activity. Upon gaining kinase activity, the receptor complex phosphorylates signaling intermediates, like STAT4. In the third step, phosphorylated signaling intermediates localize to the nucleus and promote gene transcription. Yet the different responses to IL-12 in B16F0 versus 2D6 cells suggest that the first step uses one of three competing mechanisms to create an activated receptor complex: a canonical ligand-dependent mechanism, a non-canonical receptor-dependent mechanism, and a hybrid model comprised of both canonical and non-canonical mechanisms. To select among these competing mechanisms, we used the data presented in Figure 4a-d coupled with a Bayes Factor analysis. First we stratified the flow cytometry results into different subsets based on IL12RB2 expression (see Figure S1, panel a) and quantified the phosphorylation of Akt and STAT4 in each IL12RB2 subset under different treatment conditions and in each cell line (Figure 5). We also confirmed that the observed increases in Akt and STAT4 phosphorylation with increasing IL12RB2 abundance were not attributed to fluorescent spillover (see Figure S1, panels b and c). Mathematical relationships were developed based on the three competing mechanisms and fit to the experimental data (see Methods). Bayes Factors were used to compare how well each competing mechanism captured the data. At the bottom of Figure 5, pairwise *B_ij_* comparisons were summarized in the form of a matrix, where the *i* and *j* indicate the row and column matrix entry.

**Figure 5:**
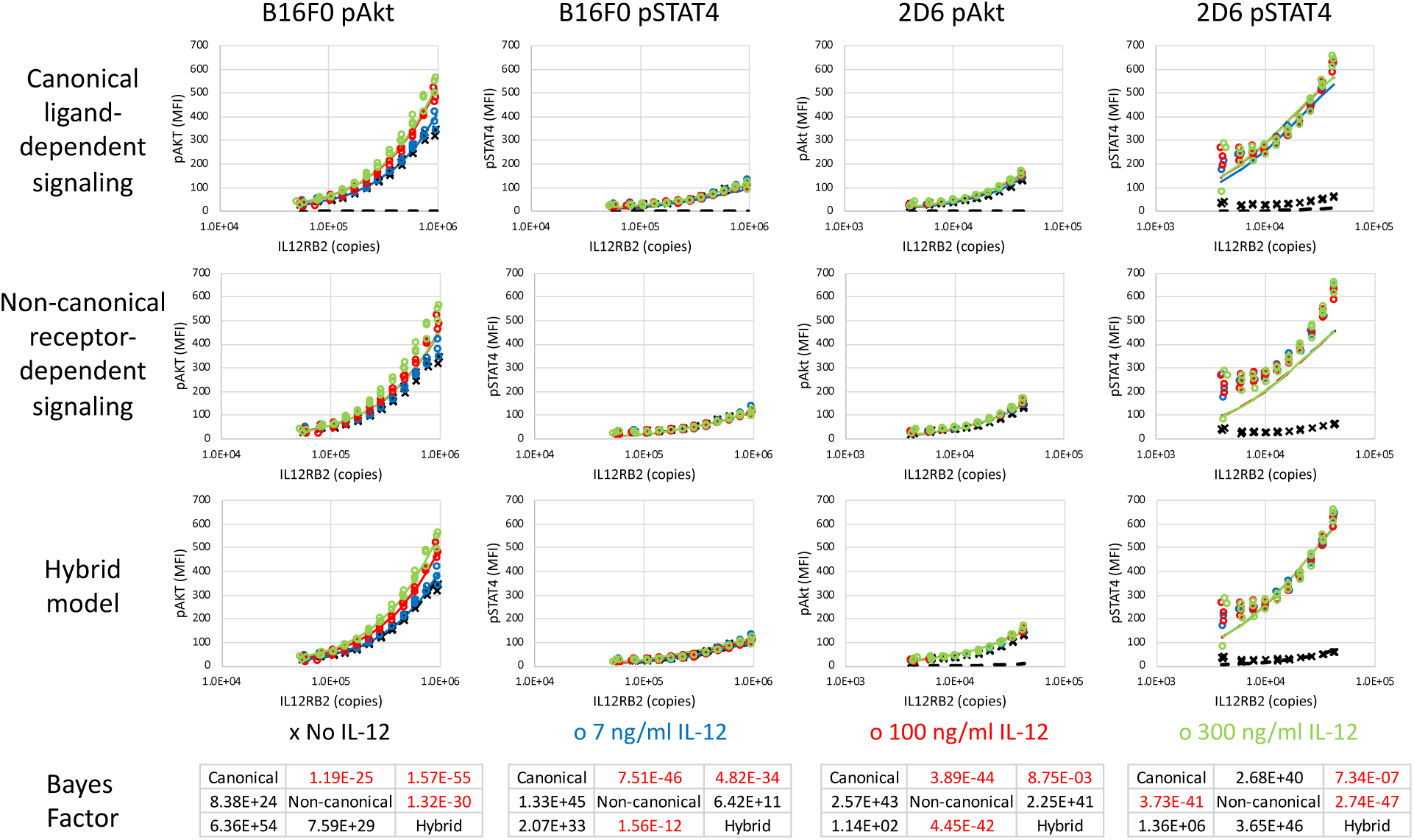
Canonical ligand-induced activation predominantly captures STAT4 activation in 2D6 cells while Akt activation in B16F0 cells is described by a hybrid model involving both non-canonical ligand-dependent and receptor-dependent activation. Stratifying flow cytometry events summarized in Figure 4a-d by IL12RB2 expression, phosphorylation of Akt and STAT4 for a subset of B16F0 (left two columns) and 2D6 (right two columns) cells are shown versus the corresponding subsets IL12RB2 abundance, expressed in copies per cell. Experimental results (symbols) are compared against simulation results using the maximum likelihood parameter estimates (lines) for three models: canonical ligand-dependent signaling (top row), non-canonical receptor-dependent signaling (second row), and a hybrid model encompassing both canonical and non-canonical models (third row). The Bayes Factor (bottom row) was calculated for each pairwise comparison between models, where *i* and *j* denote the row and column of the *B_ij_* entry. Colors indicate untreated cells (black, x) and stimulation with 7 (blue, o), 100 (red, o), and 300 (green, o) ng/mL of IL-12.

In regressing the mathematical models to the data, the ligand binding constant, *K_C_*, was determined to be 0.320 and 60.7 ng/ml in 2D6 and B16F0 cells, respectively. These values suggest that IL-12 has a 190-times higher affinity for IL12RB1:IL12RB2 compared to a receptor comprised of IL12RB2 only, as similarly reported (31). Interestingly, the canonical ligand-dependent mechanism provided the worst explanatory power in three out of four combinations. Specifically, the canonical model was unable to capture the increasing phosphorylation of Akt in both B16F0 and 2D6 cells and of STAT4 in B16F0 cells when cells with increasing copies of IL12RB2 were left unstimulated. In contrast, the non-canonical receptor-dependent mechanism provided the best explanation for phosphorylation of STAT4 in B16F0 cells and of Akt in 2D6 cells, although the extent of phosphorylation was low in both cases. In B16F0 cells, the non-canonical model captured the central trend whereby Akt phosphorylation increased with increasing IL12RB2 abundance but failed to capture IL-12-dependent variation that increased with IL12RB2 copies. For STAT4 phosphorylation in 2D6 cells, the non-canonical model provided the worst explanatory power. The hybrid model provided the best explanation of Akt phosphorylation in B16F0 cells. Moreover, knocking down IL12RB2 to below 100,000 copies in B16F0 cells is sufficient to eliminate receptor-dependent and IL-12-induced phosphorylation of Akt. For STAT4 in 2D6 cells, the hybrid model slightly edged out the canonical model as it was able to capture the slight increase in STAT4 phosphorylation observed in unstimulated cells. The high variability of STAT4 phosphorylation among replicates in 2D6 with low IL12RB2 copies is attributed to the low number of flow cytometric events included in these subsets.

Given the general trend in 2D6 cells where STAT4 and Akt phosphorylation increased with increasing IL12RB2 abundance, we next asked whether the observed responses in 2D6 cells and B16F0 cells represent different regions within a common signaling mechanism. Signals transduced by a receptor depend on assembly of the receptor complex, which can be facilitated by ligand binding. However, molecular crowding of receptors on the plasma membrane can increase ligand-independent signaling by promoting complex assembly (32). To explore the effect of molecular crowding, we plotted STAT4 and Akt phosphorylation against the estimated surface density of IL12 receptor complex (IL12R, Figure 6). For 2D6 cells, the receptor complex was assumed to be a 1:1 ratio of IL12RB1 to IL12RB2 and the complex copy number corresponded to copies of IL12RB2. For B16F0 cells, the receptor complex was assumed to be a homodimer of IL12RB2 and the complex copy number corresponded to half of the measured copies of IL12RB2. The available membrane surface area for 2D6 and B16F0 cells were estimated to be 104 and 344 *µ*m^2^, respectively (33), and adjusted based on a normalized forward scatter area (FSC-A) obtained by flow cytometry as a measure of cell size (34).

**Figure 6:**
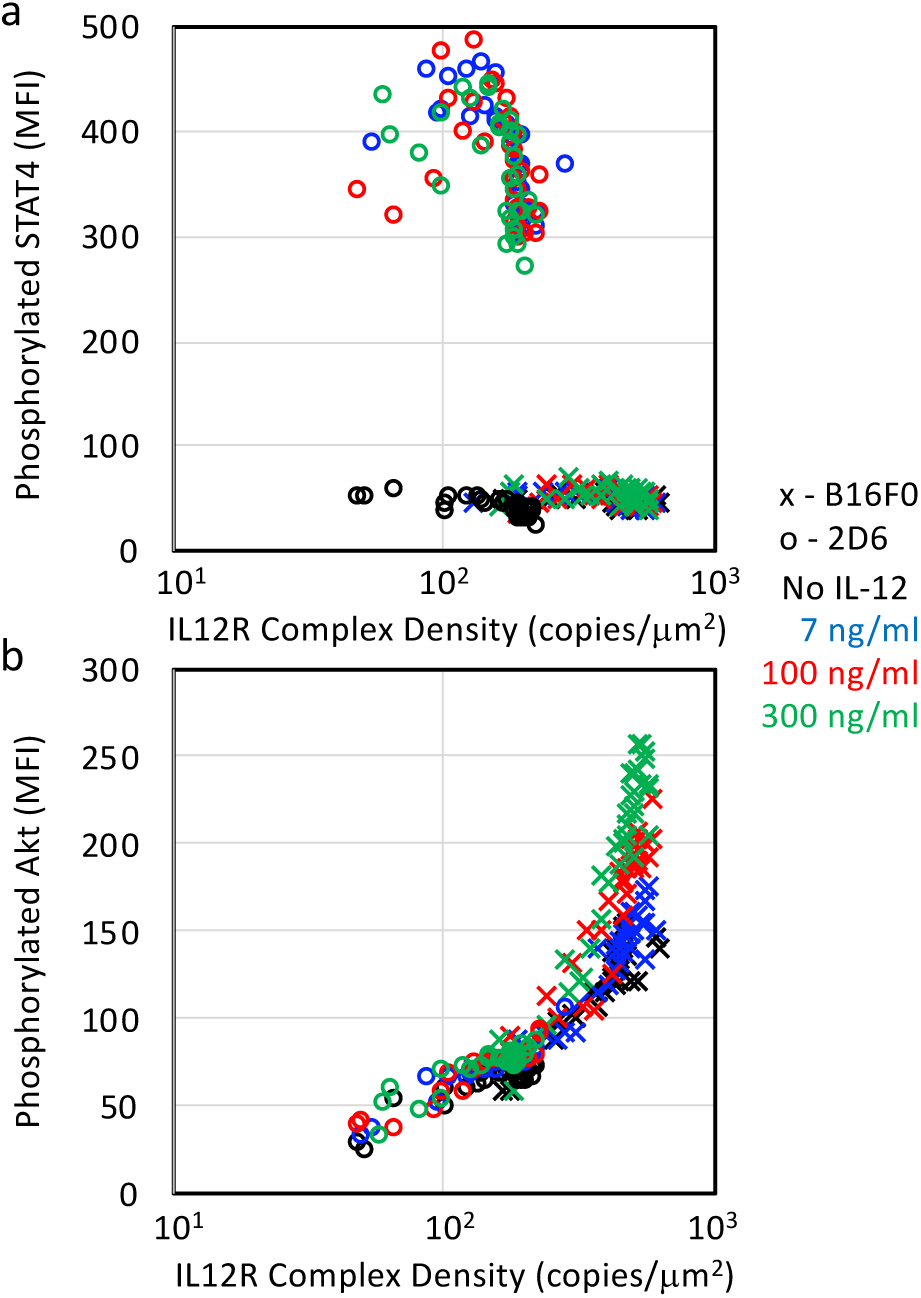
STAT4 activation depends differently on the density of IL12 receptor complexes in 2D6 cells than in B16F0 cells while Akt activation in 2D6 and B16F0 cells exhibit similar dependence on the density of IL12 receptor complexes. Phosphorylation of STAT4 (panel a) and Akt (panel b) in 2D6 (o) and B16F0 (x) cells stratified by the density of IL12 receptor complexes: IL12RB1:IL12RB2 in 2D6 cells and IL12RB2:IL12RB2 in B16F0 cells. Colors indicate untreated cells (black) and stimulation with 7 (blue), 100 (red), and 300 (green) ng/mL of IL-12. The experimental results with the highest phosphorylation levels were regressed to Eq. 12 (black lines), with the resulting equation corresponding to the maximum likelihood fit indicated.

As expected, IL-12 stimulation increased the level of STAT4 phosphorylation in 2D6 cells, with the level almost constant with increasing IL12R density (Figure 6a). Even in the absence of IL-12, STAT4 phosphorylation remained constant as the IL12R density increased, with the levels in B16F0 cells similar than those observed in unstimulated 2D6 cells. The STAT4 response suggests that a different signaling mechanism applies to STAT4 phosphorylation in 2D6 cells versus B16F0 cells. For Akt phosphorylation (Figure 6b), 2D6 and B16F0 cells provided identical levels of activation for the lower densities of IL12R, which suggests a common signaling mechanism. Only at higher densities of IL12R were differences observed upon IL-12 simulation. To rule out confounding effects that total Akt and STAT4 may increase in parallel with IL12RB2, we stimulated B16F0 cells with serum-containing media for 12 hours and assayed Akt and STAT4 phosphorylation by flow. We make the assumption that serum induced a maximal activation of Akt and STAT4 such that the levels of phosphorylated Akt and STAT4 represent the total Akt and STAT4 expression (see Figure S4, panels a and b). In subgroups stratified by IL12RB2 expression, STAT4 phosphorylation was independent of IL12R complex density. In contrast, phosphorylated Akt seemed to increase slightly with IL12R complex density. From these data, we developed a linear relationship between total Akt and IL12RB2 density (Total Akt (MFI) = 0.352 * IL12RB2 (in copies/*µm*^2^) + 382). The total Akt was then estimated for each IL12RB2 subgroup and used for the value of TOTS in equation (12). The ratio pS/TOTS still increased both as a function of IL12R complex density and IL12 (see Figure S4c), which is consistent with a hybrid model.

In comparing the responses in Figure 6, the similar response in STAT4 for all three concentrations of IL-12 in 2D6 cells while Akt phosphorylation progressively increased with increasing concentrations of IL-12 suggests that the IL12 receptor complex in 2D6 cells (i.e., IL12RB1:IL12RB2) is more efficient in phosphorylating STAT4 than either of the complexes (i.e., IL12RB1:IL12RB2 or IL12RB2:IL12RB2) in phosphorylating Akt. More importantly, the overlapping response in Akt phosphorylation between 2D6 and B16F0 cells for similar densities of IL12R suggests that the non-canonical response observed in B16F0 cells can be simply attributed to an increase in IL12RB2 copies. In addition, a gain in Akt phosphorylation is a continuous function of IL12RB2 expression. Collectively, the results suggest that IL-12 provides a survival advantage to B16F0 cells and that IL-12 induced a different cellular response in B16F0 cells compared to 2D6 T cells, whereby phosphorylation of Akt, a non-canonical signal transducer that is associated with enhanced cell survival is favored over STAT4, which is the canonical signal transducer in the IL-12 pathway.

To extend our focus beyond mouse cell models, we also wondered whether similar patterns of gene expression are observed in human melanoma based on single cell RNA-seq analysis. In a study of 31 human melanoma tumors (Figure 7), malignant melanocytes, CD4+ T cells, and NK cells expressed measurable mRNA reads for both IL12RB1 and IL12RB2 (20.7%, 21.8%, and 23.9%, respectively) while only 10.3% of CD8+ T cells had non-zero reads for both IL12RB1 and IL12RB2. The fraction of cells with non-zero reads for IL12RB1 and not IL12RB2 increased progressively in malignant melanocytes, NK cells, CD4+ T cells, and CD8+ T cells, where the proportion being 55.9%, 65.2%, 72.7%, and 81.6%, respectively. Conversely, the fraction of cells with non-zero reads for IL12RB2 and not IL12RB1 decreased from 5.2% in malignant melanocytes to 0.3% in CD8+ T cells (5.2, 2.2, 0.8, and 0.3). The distribution in cells with non-zero reads for only IL12RB1, only IL12RB2, or both were statistically different among cell types (Fisher exact Test p-value *<* 0.0005). It follows then that the ratio of IL12RB2 to IL12RB1 expression was higher for malignant melanocytes compared to both CD4+ and CD8+ T cells (p-value = 1e-5 for both). Interestingly, CD8+ T cells seemed to predominantly have an IL12RB2:IL12RB1 ratio of less than 1 (median = 0.085). The distribution in the IL12RB2:IL12RB1 ratio for NK cells seemed similar to CD4+ T cells, although there were too few cells (n=22) to assess significance. Collectively, these data suggest that human melanoma cells have an increased expression of IL12RB2 relative to IL12RB1 that is different than observed in immune cells that canonically respond to IL-12, such as NK cells, CD4+ T cells, and CD8+ T cells and qualitatively mirrors what we observed with mouse cell lines.

**Figure 7:**
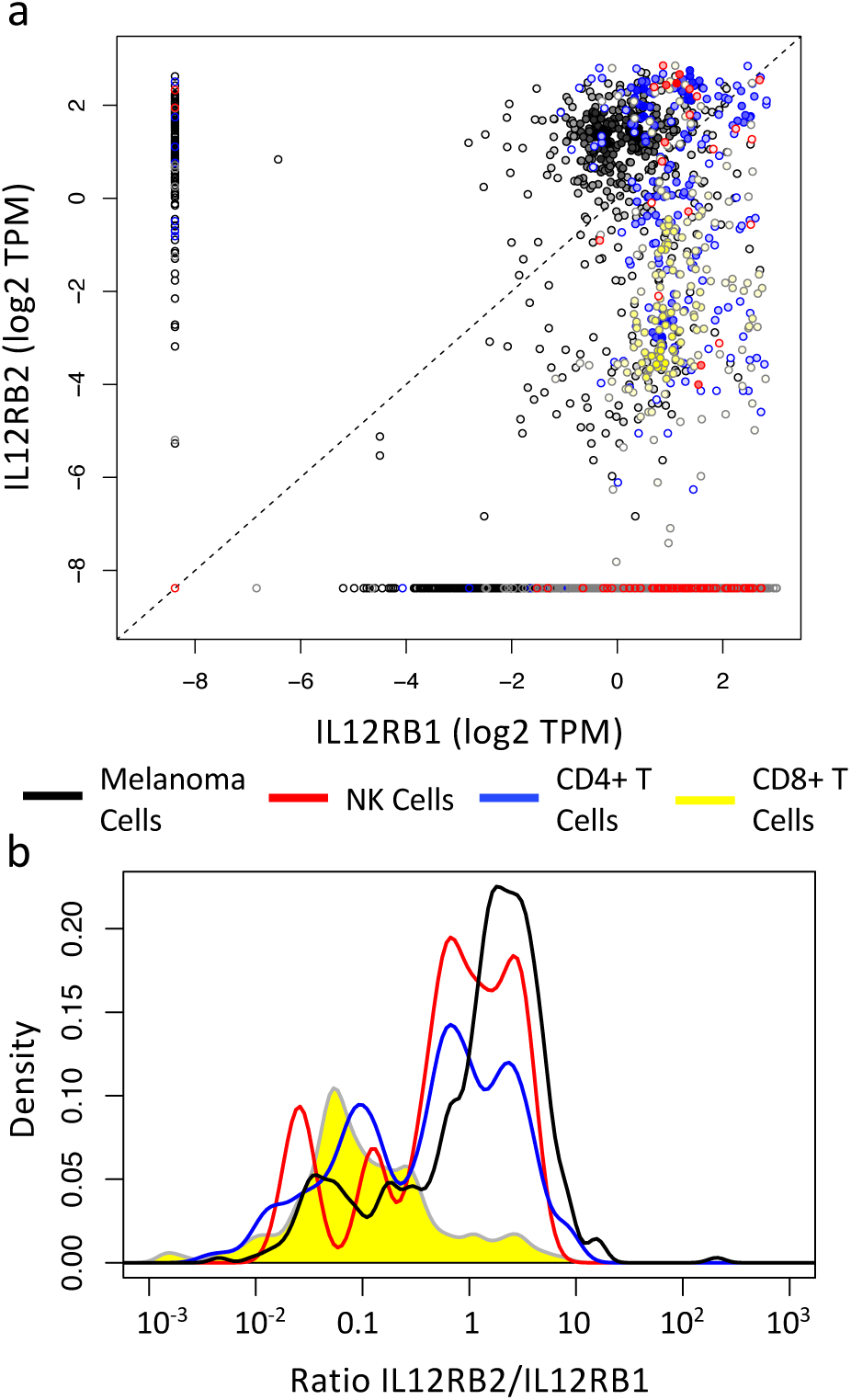
The ratio of IL12RB2 to IL12RB1 mRNA is increased in human malignant melanocytes compared to CD4+ and CD8+ T cells isolated from melanoma tumors. Single cells were isolated from 31 melanoma tumors and subject to single-cell RNA-seq analysis. (a) Scatter plot for IL12RB1 and IL12RB2 mRNA expression in TPM among malignant melanocytes (n = 2018, black), NK cells (n = 92, red), CD4+ T cells (n = 856, blue), and CD8+ T cells (n = 1759, yellow). (b) Using the subset with non-zero values for the expression of both genes, the distribution in the ratio of IL12RB2:IL12RB1 expression among malignant melanocytes (n = 418, black), NK cells (n = 22, red), CD4+ T cells (n = 187, blue), and CD8+ T cells (n = 181, yellow) is shown.

## Discussion

Given the importance of IL-12 in promoting type 1 anti-tumor immune response, we had previously observed that, by overexpressing one component of the IL-12 receptor IL12RB2, B16 melanoma cells could create a “cytokine sink” and deprive tumor-infiltrating lymphocytes of this local signal to sustain an anti-tumor response (22). Here, our results indicate that IL-12 provided an intrinsic survival advantage to B16 melanoma cells and acted directly on B16 melanoma cells to phosphorylate Akt (Figure 4a and 4c), which is in contrast to the canonical phosphorylation of STAT4 in T helper cells (Figure 4b and 4d). The survival benefit of IL-12 was mediated through IL12RB2, as knock down of IL12RB2 abrogated the effect. As the IL-12 rescue was not observed with Melan-A cells, the collective results suggest that melanoma cells rewire signal transduction to engage the non-canonical signal transducer Akt to promote survival in response to cytokines within the tumor environment.

In Figure 8, we propose a mechanism by which rewiring of the canonical IL-12 pathway could occur that is initiated by homodimerization of IL12RB2 and is enhanced by IL-12 stimulation and molecular crowding. IL-12 has been shown to promote a low but significant level of homodimerization of IL12RB2 subunits (35). Overexpression of IL12RB2 makes this association more likely to occur due to molecular crowding, as an increased number of receptors on the cell membrane increases the probability that a given receptor can get close enough to another receptor to phosphorylate the cytoplasmic domain. Following IL12RB2 homodimerization, the receptor associated Janus kinase, JAK2, interacts to promote cross phosphorylation and activation (36). Activated JAK2 can then recruit and activate PI3K via the p85a subunit (37, 38). While other intermediates may play a role, activated PI3K can then directly activate Thr308-primed Akt via phosphorylating the Ser 472/473 loci (39, 40). Our findings support the direct role of PI3K in the observed phosphorylation of Akt to affect cell survival, as inhibitors directed against PI3K pathway like LY294002 ameliorated this effect. While the data suggests that phosphorylation of Akt can occur with the IL12RB1:IL12RB2 complex, the copies of IL12RB1:IL12RB2 in 2D6 cells are too low to have a functional impact.

**Figure 8:**
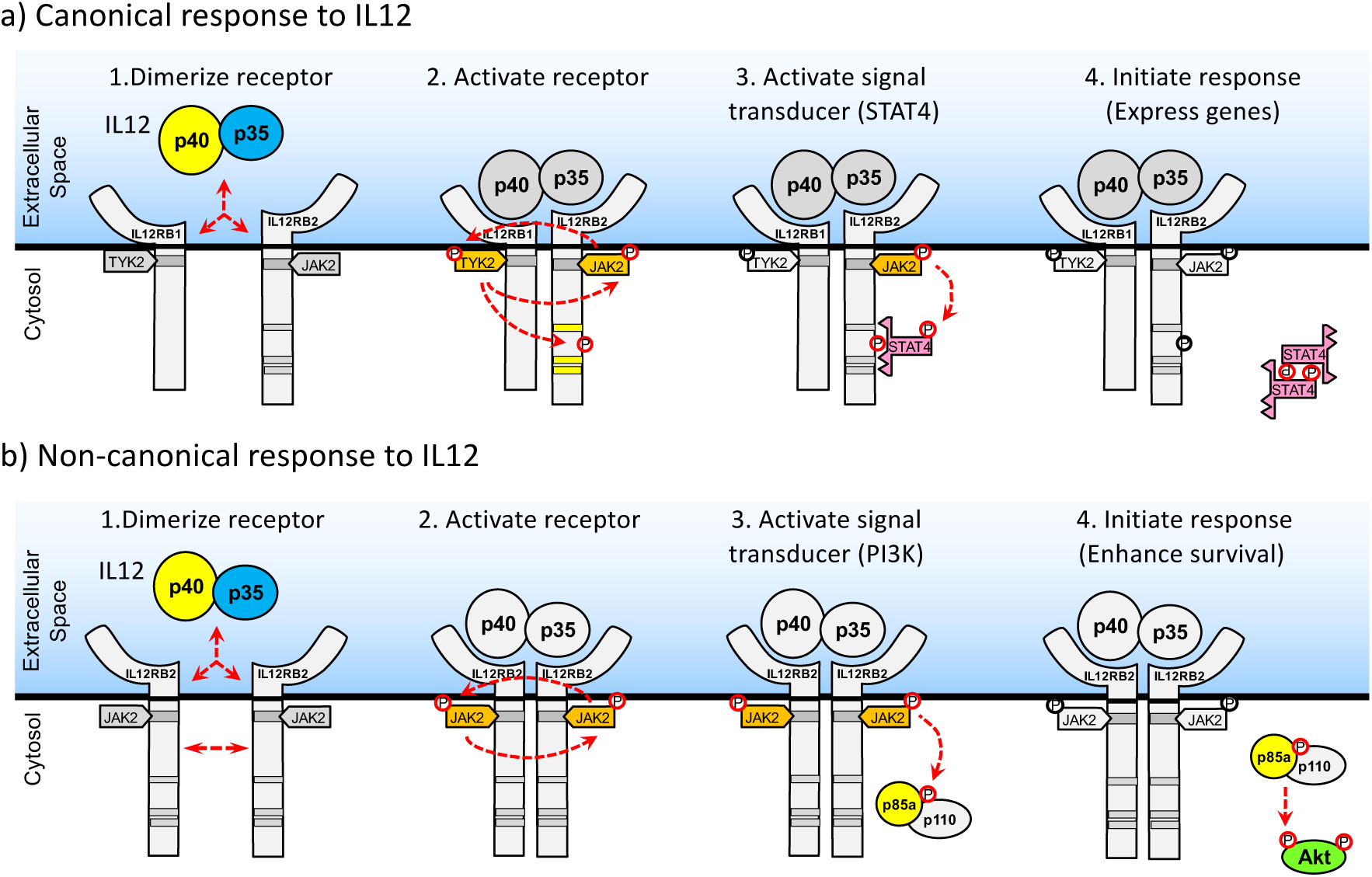
Schematic diagram illustrating the different in canonical versus non-canonical response to IL-12. (a) The canonical IL-12 pathway is triggered by (1) ligand-mediated heterodimerization of the receptor subunits IL12RB1 and IL12RB2. (2) Formation of the receptor complex promotes the phosphorylation of receptor associated kinases JAK2 and TYK2 and of a signal transducer binding site on the cytoplasmic tail of IL12RB2. (3) Phosphorylation of this IL12RB2 binding site attracts STAT4 that is subsequently phosphorylated by JAK2. (4) Phosphorylated STAT4 dimerizes and translocates into the nucleus to initiate STAT4-mediated transcription that promotes cell proliferation and Interferon-gamma production. (b) The non-canonical IL-12 pathway is triggered by (1) IL-12-enhanced homodimerization of IL12RB2 subunits. (2) Homodimerization of IL12RB2 activates the JAK2 receptor associated kinases, which in turn (3) recruit and activate PI3K via the p85a subunit. (4) Activated PI3K then phosphorylates Thr308-primed Akt at the Ser 472/473 loci, which engages the canonical Akt response that includes enhanced survival.

Rewiring of signaling pathways during oncogenesis is an important concept in cancer biology (41). For instance, a recent study across all known human kinases showed there are six types of network-attacking mutations that perturb signaling networks to promote cancer, including mutations that dysregulate phosphorylation sites to cause constitutive activation or deactivation, rewire networks through upstream or downstream interactions, and dysregulate network dynamics by activated or deactivating nodes of response (42). As for precedence of non-canonical signaling within the JAK-STAT pathway, STAT3 has been reported as a non-canonical activator of the NF-*κ*B pathway to promote myeloid-derived suppressor cells in the tumor microenvironment (43). Beside oncogenic fusion proteins, increased abundance of intracellular signaling proteins through either copy number amplifications or epigenetic changes can also create non-canonical protein interactions that could redirect the flow of intracellular information (44). Given that the Akt response between the 2D6 and B16F0 cell lines are the same where the IL12 receptor complex densities overlap, favorable changes in coding sequence are not required but merely changes that increase the abundance of IL12RB2 are sufficient to impart a fitness advantage. Moreover, the improvement in fitness, as suggested by increased Akt phosphorylation, is a continuous function of IL12RB2 copies. Knowing the basis for cancer cell rewiring can be used for therapeutic advantage. For instance, the non-canonical use of signaling cross-talk downstream of the EGF receptor sensitized resistant triple negative breast cancers to DNA-damaging chemotherapies by inhibition of EGFR (45). Similar approaches could be applied to circumvent rewiring of the IL-12 pathway.

In summary, IL-12 provides an important function in coordinating an anti-tumor immune response. In melanoma, overexpression of one component of the IL-12 receptor, IL12RB2, by malignant melanocytes may not only serve to deprive immune cells of this important cytokine, but malignant melanocytes may also use IL-12 as a means of activating cell survival pathways. Supported by prior studies, we present evidence showing that IL-12 increased the dose of imatinib chemotherapy needed to induce cell death in B16 melanoma cells and that IL-12 activated a non-canonical cell survival protein, Akt, in a melanoma cell line, but not in the 2D6 T helper cell line. While subsequent studies will help clarify the mechanistic details to improve the translational impact, these studies highlight the importance of non-canonical pathway activation in cancer.

## Author Contributions

CNBH: planned experiments, performed experiments, analyzed data, and drafted manuscript; WD: planned experiments and performed experiments; OM: performed experiments; PW: performed experiments; YR: planned experiments; DJK: planned experiments, analyzed data, performed quantitative analysis, and drafted manuscript. All authors edited and approved of the final version of the manuscript.

## Acknowledgments

The authors thank Cassidy Bland and Adam Palmer for assisting with manuscript preparation and Vanessa Cuppett and Ning Cheng for technical assistance. Sources of funding that contributed to this work include NIH/NCI R01CA193473, NSF 1644932, WVU Flow Cytometry Core funding: MBRCC CoBRE grant GM103488/RR032138, ARIA S10 grant RR020866, Fortessa S10 grant OD016165, and WV InBRE grant GM103434. The content is solely the responsibility of the authors and does not necessarily represent the official views of the National Cancer Institute, the National Institutes of Health, or the National Science Foundation. CBH, WD, OM, PW, YR, and DJK declare that they have no competing interests.

---

**Supplemental Figure S1:**
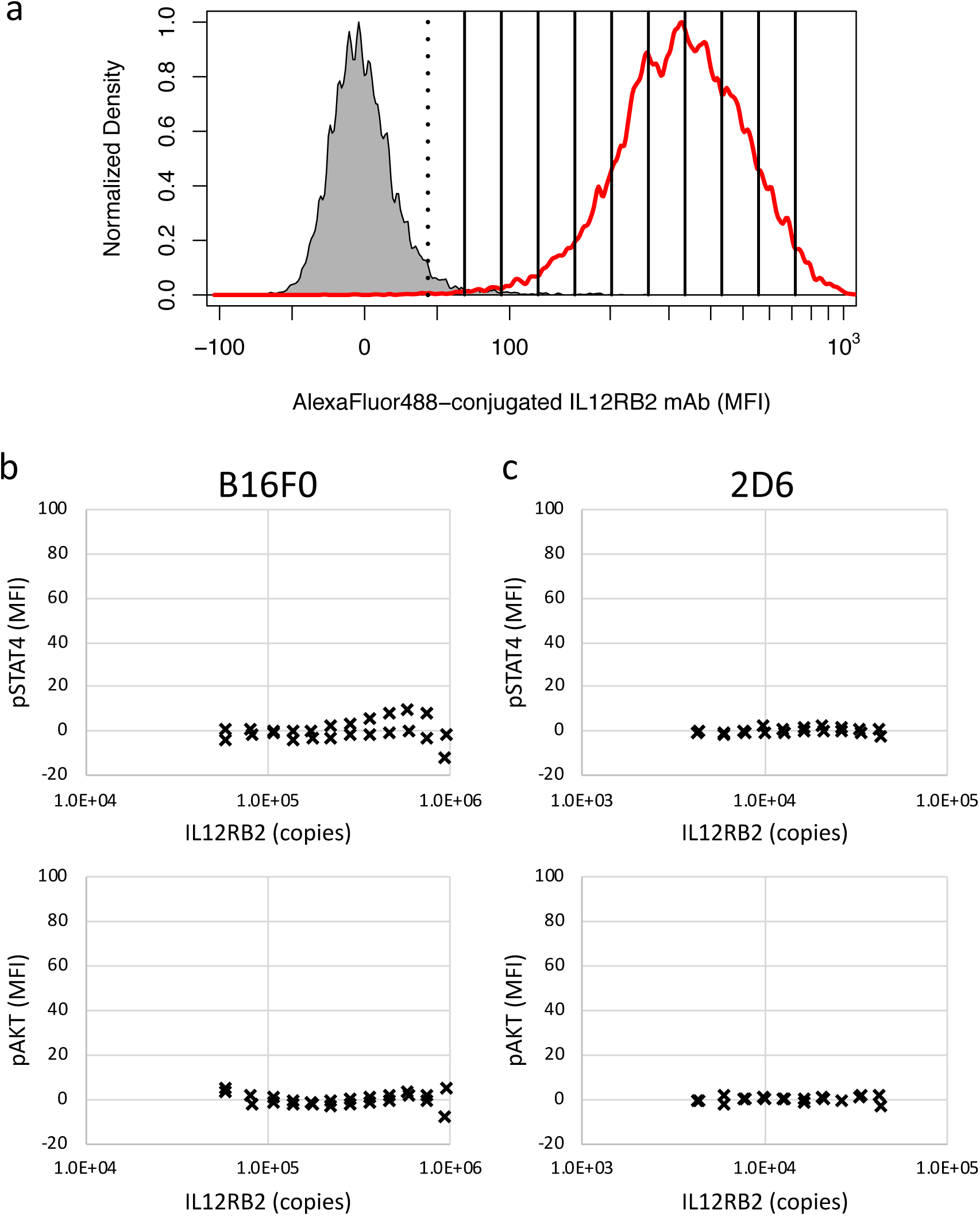
Cell subsets were established that vary in expression of IL12RB2 and with minimal fluorescent spillover into pAkt and pSTAT4 measures. (a) Flow cytometric events were subdivided into groups based on IL12RB2 expression, where solid vertical lines denote limits of IL12RB2 expression associated with each subgroup. Red curve denotes distribution in IL12RB2 expression by B16F0 while gray shaded indicates unstained controls. Single-stained controls for IL12RB2 in B16F0 (b) and 2D6 (c) cells were used to establish that fluorescence associated with measuring Akt and STAT4 phosphorylation was not due to fluorescent spillover. Results for two independent experiments are shown for each subgroup based on IL12RB2 expression.

**Supplemental Figure S2:**
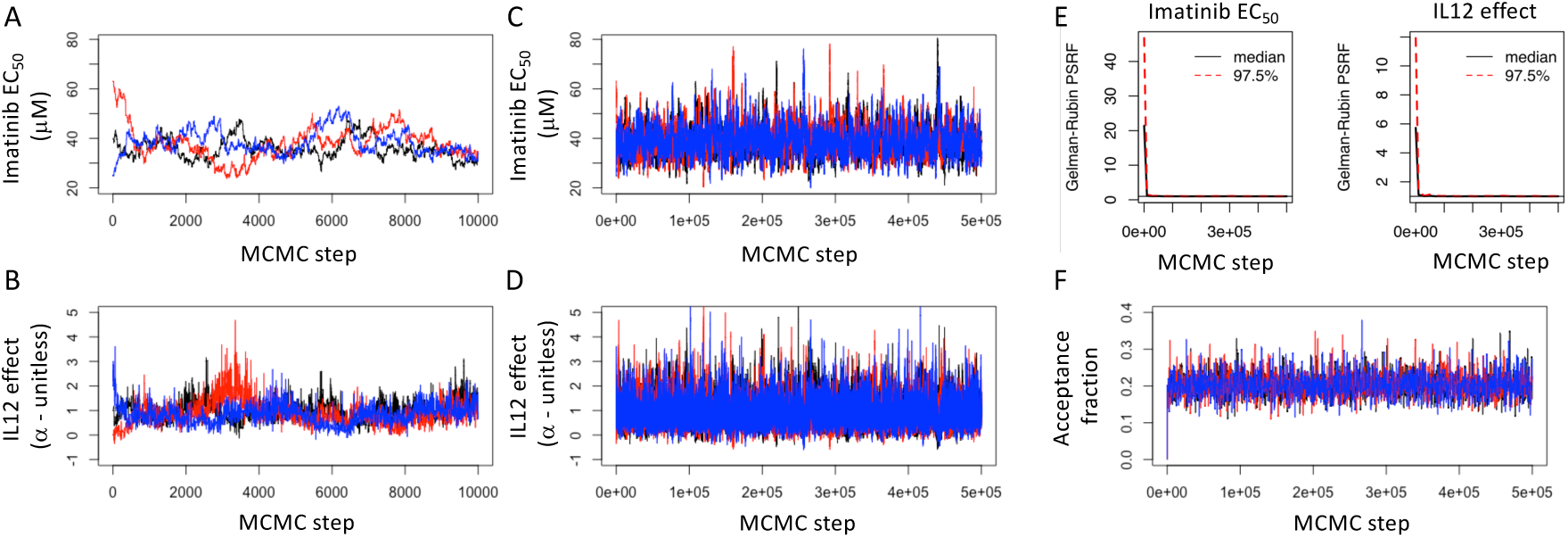
Diagnostics for Markov Chain Monte Carlo estimates of the posterior distribution in the Imatinib *EC*_50_ and IL12 effect (*α*) estimated from flow cytometry viability assay using B16F0 cells. The traces for the initial 10000 steps (A and B) and full length (C and D) of three independent chains (colored in black, blue, and red, respectively) with over disperse starting points. (E) The Gelman-Rubin potential scale reduction factor (PSRF) was used to assess convergence of the Markov Chains to the posterior distribution in the Imatinib *EC*_50_ (left panel) and IL12 effect *α* (right panel), where a value of less than 1.2 indicated that the chains have converged to sampling the posterior distribution. The MCMC chains converged after less than 50,000 steps. (F) New steps in the Markov Chain were proposed using a normally distributed random number generator with a mean of zero and adjusted standard deviation such that the acceptance fraction was 0.2.

**Supplemental Figure S3:**
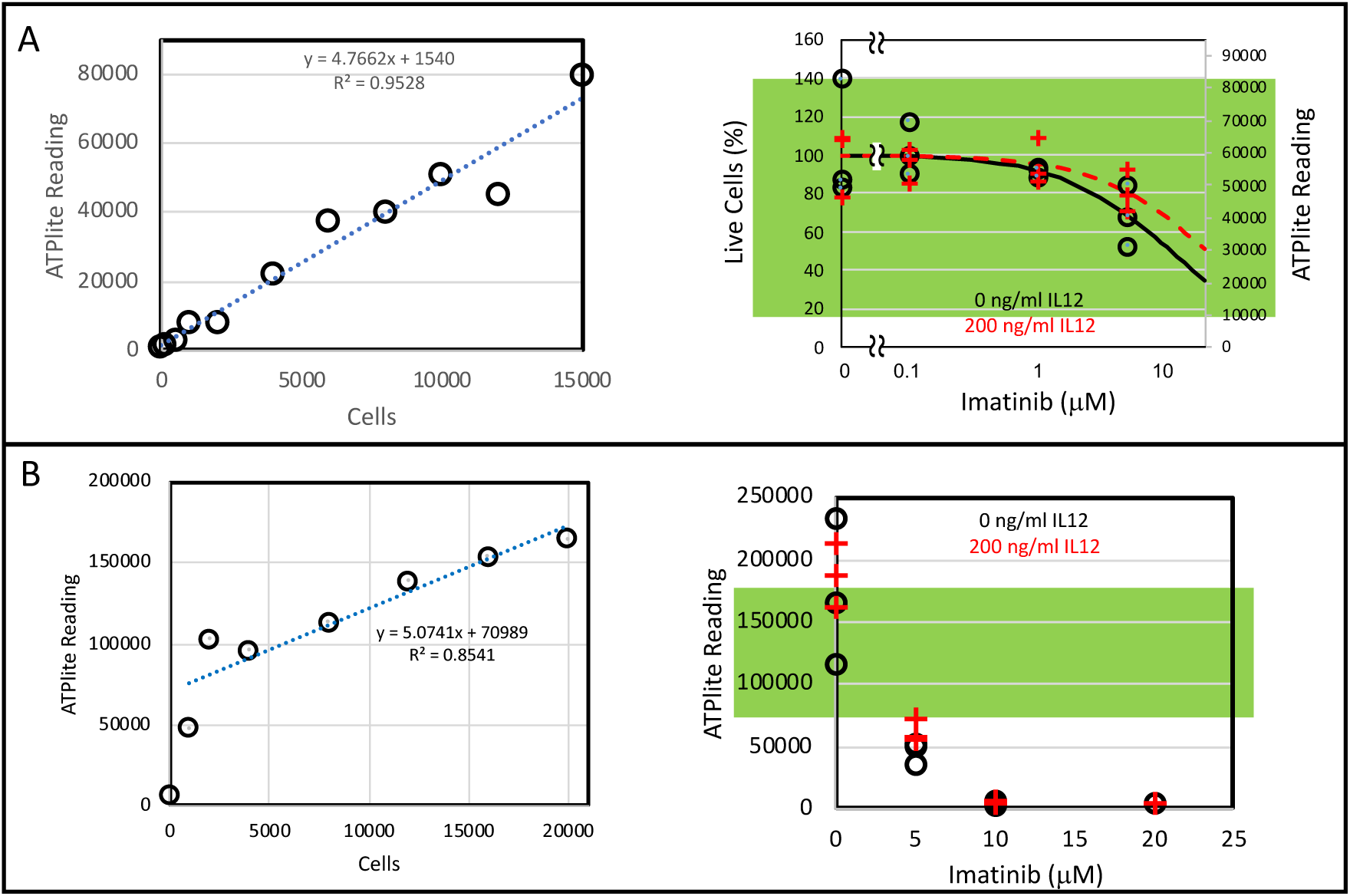
Calibration curves for quantifying cell viability using the ATPlite assay. (A) Increasing concentrations of B16F0 cells were plated just prior to reading viability using the ATPlite assay to establish the dynamic range of the assay (left panel). Results for experimental conditions that were acquired within the dynamic range of the assay are indicated by green overlay. (B) In a separate dose-finding experiment, increasing concentrations of B16F0 cells were plated just prior to reading viability using the ATPlite assay (left panel). While higher doses of imatinib appeared to reduce cell viability to near zero, the experimental conditions were acquired outside of the dynamic range of the assay (green overlay).

**Supplemental Figure S4:**
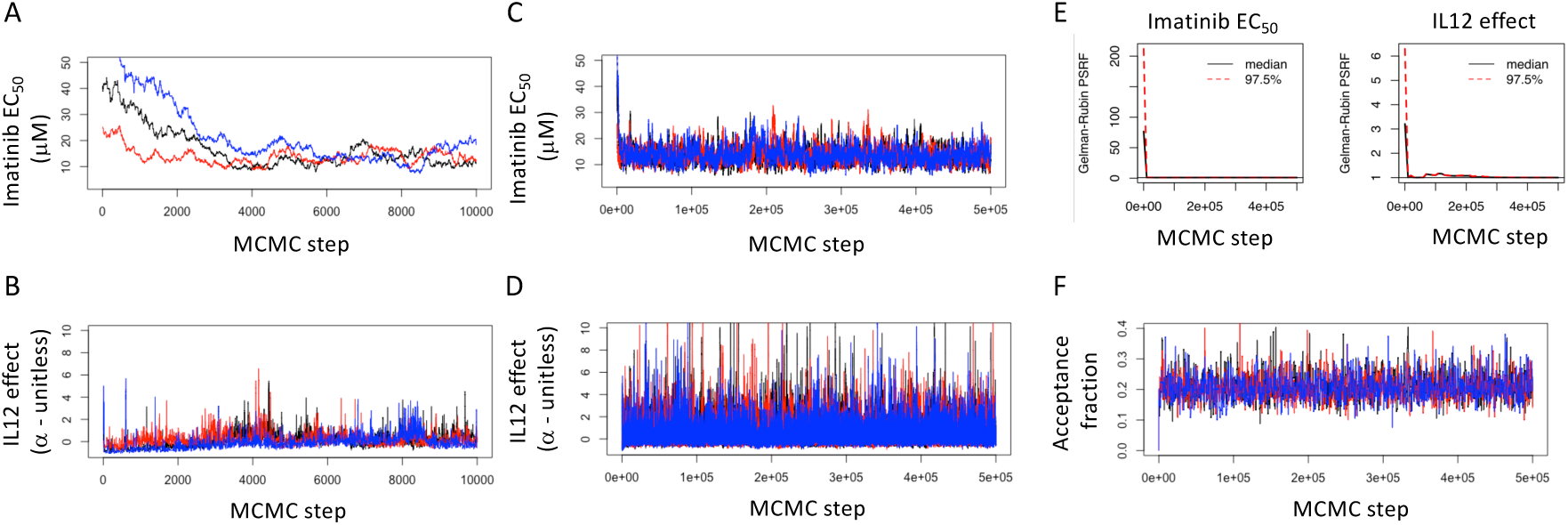
Diagnostics for Markov Chain Monte Carlo estimates of the posterior distribution in the Imatinib *EC*_50_ and IL12 effect (*α*) estimated from ATPlite assay results using Melan-A cells. The traces for the initial 10000 steps (A and B) and full length (C and D) of three independent chains (colored in black, blue, and red, respectively) with over disperse starting points. (E) The Gelman-Rubin potential scale reduction factor (PSRF) was used to assess convergence of the Markov Chains to the posterior distribution in the Imatinib *EC*_50_ (left panel) and IL12 effect *α* (right panel), where a value of less than 1.2 indicated that the chains have converged to sampling the posterior distribution. The MCMC chains converged after less than 50,000 steps. (F) New steps in the Markov Chain were proposed using a normally distributed random number generator with a mean of zero and adjusted standard deviation such that the acceptance fraction was 0.2.

**Supplemental Figure S5:**
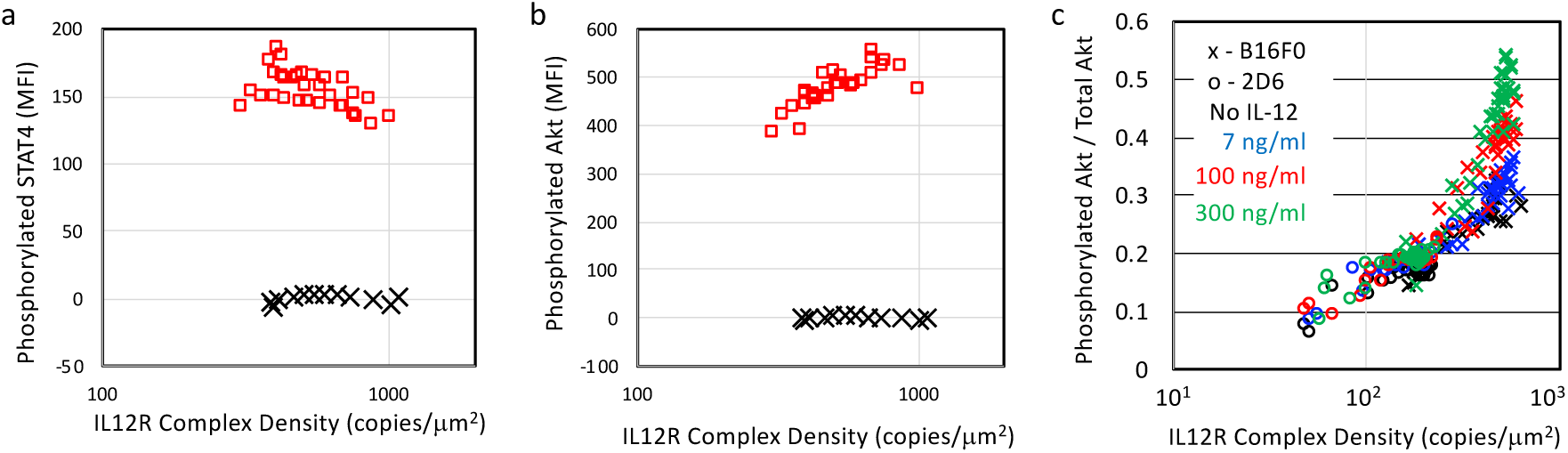
Estimating total Akt and STAT4 values. Media contained serum was used to elicit a near maximal phosphorylation of STAT4 (A) and Akt (B) in B16F0 cells following a 12 hour stimulation (red squares). Phosphorylation of STAT4 and Akt was assayed by flow cytometry, where results are shown for each subgroup based on IL12RB2 expression. Single-stained controls for IL12RB2 in B16F0 (black x?s) cells were used to establish that fluorescence associated with measuring Akt and STAT4 phosphorylation was not due to fluorescent spillover. From these data, we developed a linear relationship between total Akt and IL12RB2 density (Total Akt (MFI) = 0.352 * IL12RB2 (in copies/*µm*^2^) + 382). The total Akt was then estimated for each IL12RB2 subgroup and used for the value of TOTS in equation (12). (C) The ratio of phosphorylated Akt to total Akt, which corresponds to pS/TOTS in equations 12, 14, 16, and 19, was calculated for the different experimental conditions. Total Akt was assumed to follow the same dependence on IL12RB2 in 2D6 and B16F0 cells.

